# Taxonomic Diversity of Ants (Formicidae) in Forest Fragments of Tropical Dry Forest in Atlántico, Colombia

**DOI:** 10.1101/2025.08.13.670019

**Authors:** Damaris Grandas-Gaona, Daniel Posada-Echeverría, Valeria Machacón, Rodrigo Sarmiento, Neis Martínez, Rafik Neme

## Abstract

This study provides the first taxonomic diversity analysis of ants in the department of Atlántico, Colombia, documenting 6.75% of the country’s known ant species, 29.5% of its genera, and 63.3% of its subfamilies. We report the first record of *Leptogenys volcanica* in the Colombian Caribbean and expand the known distribution of 19 species within Atlántico. Our results show marked differences in species composition among forest fragments. El Morro exhibited the highest taxonomic distinctness (TD), supporting a community composed of more distantly related species. In contrast, Uninorte, an isolated urban fragment, had the lowest TD value, suggesting a more taxonomically clustered community and potentially acting as a diversity sink. Nesting values indicate that El Morro functions as a biodiversity reservoir, while the species composition of Los Charcones appears shaped by past land-use changes. The dominance of Myrmicinae and Formicinae aligns with patterns observed in other tropical dry forests. Although fragmentation was not directly assessed, patterns of alpha, beta, and taxonomic distinctness suggest that isolation and land use may influence community composition. El Morro emerges as a key fragment for regional conservation. Enhancing habitat connectivity could help mitigate biodiversity loss in these fragmented landscapes. This study provides a critical baseline for future research on ant ecology and conservation in the Caribbean region of Colombia.

## 2. Introduction

Ants are vital components of ecosystems, playing key roles in nutrient cycling, seed dispersal, and pest control. Their high diversity and ecological sensitivity make them valuable bioindicators of environmental health and ecosystem change (1). With over 15,000 described species worldwide (2),ants are especially diverse in tropical regions, including Colombia— one of the world’s megadiverse countries (3). However, habitat fragmentation and human activities are major threats to this diversity, particularly in tropical dry forests (TDFs) (4).

TDFs are among Colombia’s most endangered ecosystems. Once covering 8.88 million hectares, they have been reduced to just 1 million hectares (5,6). The Caribbean region, which contains 40.9% of Colombia’s TDFs, has lost over 94% of its original extent, with only 5.8% remaining as mature forest due to agricultural expansion and urbanization (7). These disturbances severely impact ant communities, leading to declines in species diversity and ecological functionality (8,9).

Despite the critical ecological roles of ants, research on their diversity in Colombia’s dry forests remains limited, especially in Atlántico department. While national datasets exist (10), only three studies in the past decade have examined ant diversity in this region (11–13). These studies focused primarily on two subfamilies (Ponerinae and Ectatomminae), leaving significant gaps in knowledge about key genera such as *Pseudomyrmex* Lund, 1831, *Camponotus* Mayr, 1861, *Cephalotes* Latreille, 1802, and *Pheidole* Westwood, 1839.

To address the lack of information on ant communities in the tropical dry forests of the Atlántico department, this study documents their diversity across forest fragments with varying degrees of disturbance. By analyzing species composition, distribution, and phylogenetic relationships, we aim to deepen the understanding of the region’s myrmecofauna, evaluate their potential as bioindicators, and inform conservation strategies. We hypothesize that habitat fragmentation and different land-use types affect not only species richness and composition, but also phylogenetic diversity, with better-preserved fragments supporting more diverse and representative evolutionary lineages than urban or heavily altered remnants.

## 3. Methods

### Study Area

The Atlántico department, located in northwestern Colombia (10°16’27”–11°06’52” N, 74°42’54”–75°17’13” W), covers 3,470 km^2^ and includes remnants of tropical dry forests (TDF). These forests experience pronounced seasonality, with a dry period from December to May. The region’s climate is warm, with an average annual temperature of 27.8–29°C, relative humidity between 60% and 90%, and annual precipitation ranging from 926 to 1,000 mm (7).

The TDFs from Departamento del Atlántico correspond to a hygrotropophytic formation of tropical dry forest (TDF) characteristic of the Colombian Caribbean region (14). The study area generally exhibits a bimodal, four-season rainfall regime, with two main rainy seasons occurring from April to June and from August to November. The average annual precipitation is 1260 ± 347 mm, with mean annual temperatures of 27.3 ± 0.9 °C (7).

This study was conducted in five TDF fragments in Atlántico, each characterized by vegetation from families such as Malvaceae, Fabaceae, Anacardiaceae, Cactaceae, Lecythidaceae, Sapindaceae, Euphorbiaceae, and Polygonaceae. Sites were selected based on their accessibility and availability following the 2020 pandemic, when mobility restrictions were still in place. Forest areas of varying sizes and levels of disturbance were prioritized to capture greater ecological variability **Table 1**.

**Table 1.**
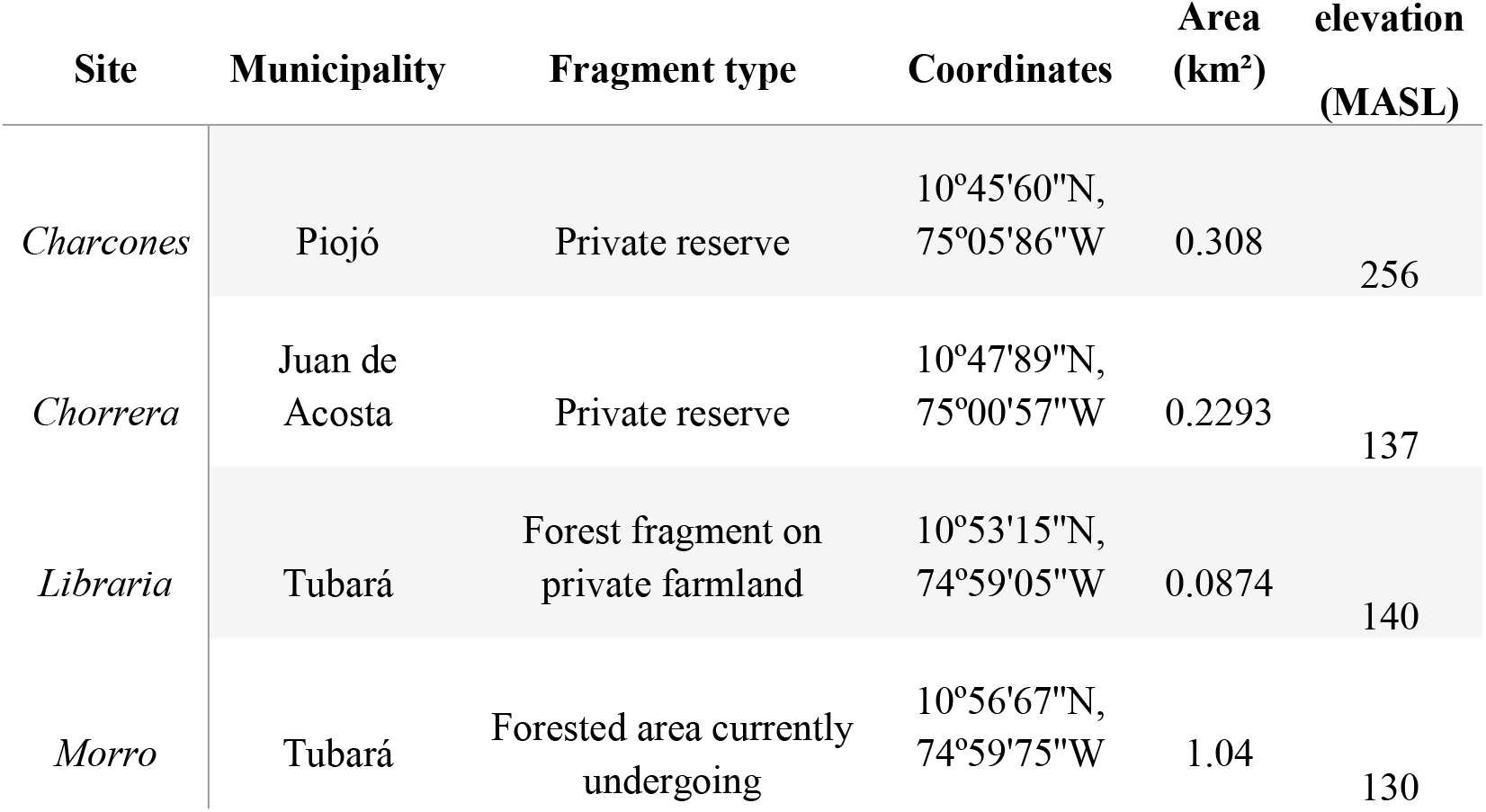
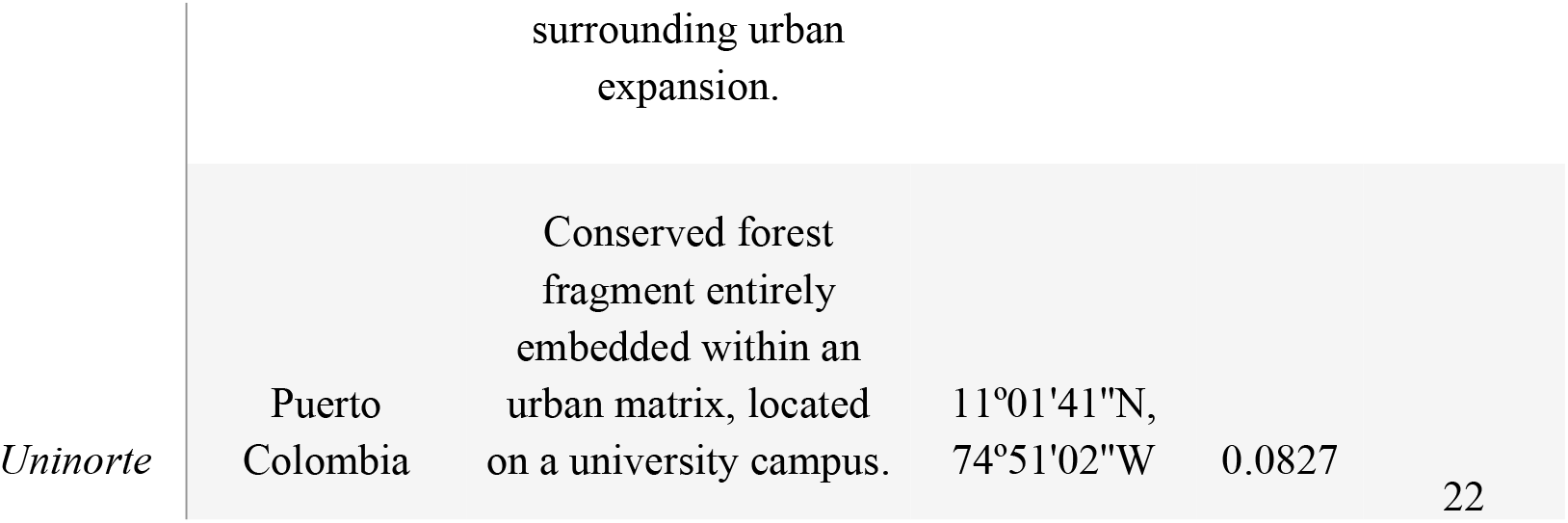
Area Tropical dry forest fragments selected for the study in the department of Atlántico, Colombia. The analyzed fragments vary in their degree of anthropogenic disturbance, including private reserves, urban remnants, and forests surrounded by agricultural matrices.

### Sampling Methods

Sampling was conducted from July to October 2021 using a standardized protocol to ensure comparability across sites. In each forest fragment, a 200-meter transect was established, with five equidistant sampling stations. At each station, ants were collected using four complementary methods:

- Pitfall traps – Two traps per station, left active for seven days.
- Manual collection – A team of three people conducted a 10-minute manual collection within a 4-meter radius around each pitfall trap.
- Baited traps – Three Cornell traps baited with tuna and sugar, deployed for three hours. The first trap was buried 10 centimeters deep in the soil, the second was placed on the leaf litter, and the third was installed in the trees at a height of 1.50 meter.
- Malaise traps – One trap per station, left active for seven days.

Malaise traps—though not typically used for ant surveys—have demonstrated strong effectiveness in sampling arboreal and flying insects. A study conducted in an urban tropical dry forest in Puerto Colombia showed that Malaise traps captured the highest richness of insect morphospecies compared to pitfall and fogging traps, and contributed many unique taxa, highlighting the complementarity of these methods in TDF sampling(15). Winkler traps were not used due to the unavailability of the equipment at the time of sampling.

All collected specimens were preserved in 96% ethanol and processed at Universidad del Norte. Identification was performed using taxonomic keys from Fernandez *et al*. (10).

Sampling done under the Framework Permit for the Collection of Specimens of Wild Species of Biological Diversity for Non-Commercial Scientific Research Purposes, 0739 from July 8th, 2014 valid through 2024, granted to Universidad del Norte by the Colombian National Authority of Environmental Licenses (ANLA).

### Data Analysis

To assess ant diversity in Atlántico, we compiled species records from scientific publications, environmental authority reports (16), and public databases (17,18). These records were compared with the species collected in this study.

For species counts, the maximum number of recorded species per genus across different sources was used as a reference. If one study reported 10 species and another reported 12, the count of 12 was used. Morphospecies were assigned species names, when possible, otherwise recorded at the genus level as Genus sp. to prevent overestimation of species richness.

To evaluate sampling completeness, rarefaction and extrapolation curves of Hill numbers were generated using the iNEXT package in R (19), based on frequency data (20). To compare species richness across fragments, we calculated the diversity index (Chao et al., 2020). Taxonomic diversity was assessed using the average taxonomic distinction (TD), which measures differentiation within communities, this analysis provides an approximation of the phylogenetic diversity of the study sites. (21). These analyses were based on a presence/absence matrix incorporating subfamily, genus, and species categories. Taxonomic diversity calculations were performed using the *vegan* package (*taxa2dist* and *taxondive* functions) in R (22).

The dissimilarity analysis of community composition followed the methodology of (23). Data were processed in RStudio 2022.02 using the *betapart* package, using the functions *betapart*.*core, beta*.*multi*, and *beta*.*pair*, based on the Jaccard index, which assigns equal weight to all species (24,25). These functions allowed us to assess the contribution of species turnover and nestedness in the study sites.

## 4. Results

### Ant Diversity and Composition

A total of 24,257 individual ants were collected, representing seven subfamilies, 31 genera, and 81 morphospecies (**Table 2; Table 3**). Of these, 56 species were identified at the species level, while 25 were classified as sp. The most diverse subfamilies were Myrmicinae (36 morphospecies), Formicinae (15 morphospecies) and Pseudomyrmecinae (12 morphospecies).

**Table 2.** Taxonomic Diversity, Taxonomic Distinctness (TD), Sampling Completeness. The studied sites show significant variations in species richness and taxonomic distinctness. El Morro stands out as the most diverse site across the different taxonomic categories, but it is not significantly different from other forests in terms of phylogenetic diversity (TD). Despite their high richness, Uninorte and Chorrera have low taxonomic distinctness, reflecting homogeneous communities dominated by few species. Most sites show high sampling completeness (>92%), except for Los Charcones, suggesting that the detected ant community in this site is incomplete.

| Sites | Species | Genera | Subfamilies | TD | Completeness |
| --- | --- | --- | --- | --- | --- |
| Charcones | 31 | 14 | 6 | 86,60 | 70,19% |
| Chorrera | 43 | 19 | 7 | 83,55 | 92,34% |
| Libraria | 38 | 15 | 6 | 87,59 | 87,75% |
| Morro | 62 | 24 | 7 | 87,99 | 85,53% |
| Uninorte | 41 | 20 | 6 | 83,41 | 92,08% |

**Table 3.** Taxonomic Richness of Ants at the Study Sites, Including Hierarchical Categories of Subfamily, Genus, and Species. Seven subfamilies, 31 genera, and 81 species or morphospecies were identified. The most representative subfamily was Myrmicinae, with the genera Cephalotes and Pheidole having the highest number of species, followed by Formicinae, with *Camponotus* as the predominant genus, and Pseudomyrmecinae represented solely by *Pseudomyrmex*.

| Subfamilies | Genera | Species | Charcones | Chorrera | Libraria | Morro | Uninorte |
| --- | --- | --- | --- | --- | --- | --- | --- |
| Dolichoderinae | *Azteca* | *Azteca forelii* | 0 | 0,6 | 0 | 1 | 0 |
|  | *Dorymyrmex* | *Dorymyrmex bicolor* | 0 | 0 | 0 | 0,6 | 0 |
|  |  | *Dorymyrmex biconis* | 0 | 0 | 0 | 0 | 0,4 |
|  | *Forelius* | *Forelius damiani* | 0 | 0 | 0,2 | 0 | 0 |
|  | *Tapinoma* | *Tapinoma melanocephalum* | 0 | 0 | 0 | 0,2 | 0 |
| Dorylinae | *Labidus* | *Labidus coecus* | 1 | 0,2 | 0 | 0,6 | 0 |
| Ectatomminae | *Ectatomma* | *Ectatomma ruidum* | 1 | 1 | 1 | 1 | 1 |
|  |  | *Ectatomma tuberculatum* | 0 | 0 | 0 | 0,6 | 0 |
|  | *Gnamptogenys* | *Gnamptogenys striatula* | 0,2 | 0 | 0 | 0 | 0 |
| Formicinae | *Brachymyrmex* | *Brachymyrmex* sp1 | 0 | 0 | 0,2 | 0,2 | 0 |
|  | *Nylanderia* | *Nylanderia* sp1 | 0 | 0 | 0 | 0 | 0,2 |
|  | *Camponotus* | *Camponotus substitutus* | 0,2 | 0,6 | 1 | 0,8 | 1 |
|  |  | *Camponotus dalmasi* | 0,2 | 0 | 0 | 0 | 0 |
|  |  | *Camponotus lindigi* | 1 | 1 | 1 | 0,2 | 0,6 |
|  |  | *Camponotus* sp10 | 0 | 0 | 0 | 0 | 0,2 |
|  |  | *Camponotus indianus* | 0 | 0 | 0 | 0,2 | 0 |
|  |  | *Camponotus bidens* | 0,2 | 0 | 0,6 | 0,4 | 0,6 |
|  |  | *Camponotus linnaei* | 0 | 0 | 0 | 0,6 | 0 |
|  |  | *Camponotus blandus* | 0,2 | 0 | 0,2 | 0 | 0 |
|  |  | *Camponotus* sp4 | 0 | 0 | 0,4 | 0,2 | 0,2 |
|  |  | *Camponotus* sp5 | 0 | 0 | 0,2 | 0,2 | 0 |
|  |  | *Camponotus coruscus* | 0,8 | 0,4 | 0,6 | 0,6 | 0,2 |
|  |  | *Camponotus* sp7 | 0 | 0,2 | 0 | 0,4 | 0 |
|  |  | *Camponotus* sp9 | 0 | 0 | 0,2 | 0,6 | 0 |
| Myrmicinae | *Arcromyrmex* | *Acromyrmex octospinosus* | 0 | 0,8 | 0 | 0,2 | 0,6 |
|  | *Cephalotes* | *Cephalotes atratus* | 0 | 0,2 | 0 | 0 | 0 |
|  |  | *Cephalotes columbicus* | 0,2 | 1 | 1 | 0,8 | 1 |
|  |  | *Cephalotes femoralis* | 0 | 0,2 | 0 | 0,4 | 0,4 |
|  |  | *Cephalotes maculatus* | 0 | 0,4 | 0 | 0,2 | 0,2 |
|  |  | *Cephalotes minutus* | 0 | 0,8 | 0,4 | 0,6 | 0,6 |
|  |  | *Cephalotes mompox* | 0,4 | 0,6 | 1 | 0,8 | 0,4 |
|  |  | *Cephalotes pallens* | 0 | 0,6 | 0,6 | 0,4 | 0,8 |
|  |  | *Cephalotes pusillus* | 0,2 | 0 | 0 | 0 | 0 |
|  |  | *Cephalotes* sp8 | 0 | 0 | 0 | 0 | 0,2 |
|  | *Crematogaster* | *Crematogaster distans* | 1 | 1 | 0 | 0,2 | 0,4 |
|  |  | *Crematogaster ampla* | 1 | 0,4 | 1 | 0,8 | 0,8 |
|  |  | *Crematogaster* sp3 | 0,2 | 0,4 | 0,4 | 0,4 | 0 |
|  |  | *Crematogaster crinosa* | 0,2 | 0 | 0,2 | 0 | 0,2 |
|  |  | *Crematogaster carinata* | 0 | 0,8 | 0,4 | 0,2 | 0,2 |
|  | *Cyphomyrmex* | *Cyphomyrmex rimosus* | 0,4 | 0,4 | 0,2 | 0,2 | 0,6 |
|  | *Monomorium* | *Monomorium* sp1 | 0 | 0,2 | 0 | 0,2 | 0 |
|  | *Mycetomoellerius* | *Mycetomoellerius uruchii* | 0,8 | 1 | 0,6 | 0,6 | 0,4 |
|  | *Paratrachymyrmex* | *Paratrachymyrmex irmgardae* | 0 | 0,4 | 0 | 0 | 0,8 |
|  | *Pheidole* | *Pheidole* sp1 | 0,2 | 0,2 | 0 | 0,4 | 0,4 |
|  |  | *Pheidole* sp2 | 0,2 | 0,2 | 0,2 | 0,2 | 0 |
|  |  | *Pheidole* sp3 | 0,2 | 0,6 | 0,6 | 1 | 1 |
|  |  | *Pheidole* sp4 | 0 | 0 | 0 | 0,2 | 0 |
|  |  | *Pheidole* sp5 | 0 | 0 | 0 | 0,2 | 0 |
|  |  | *Pheidole* sp6 | 0 | 0,6 | 0 | 0,6 | 0 |
|  |  | *Pheidole* sp7 | 0 | 0,6 | 0,2 | 0,6 | 0 |
|  |  | *Pheidole transversostriata* | 0,2 | 0,4 | 0,6 | 0,4 | 0,4 |
|  | *Pogonomyrmex* | *Pogonomyrmex mayri* | 0 | 1 | 0 | 0 | 0,6 |
|  | *Rogeria* | *Rogeria foreli* | 0 | 0 | 0 | 0 | 0,4 |
|  |  | *Rogeria belti* | 0 | 0 | 0 | 0 | 0,2 |
|  | *Sericomyrmex* | *Sericomyrmex opacus* | 0 | 0 | 0 | 0,2 | 0 |
|  | *Solenopsis* | *Solenopsis geminata* | 0,4 | 0,8 | 1 | 0,8 | 0,8 |
|  |  | *Solenopsis* sp2 | 0 | 0,6 | 0 | 0 | 0 |
|  |  | *Solenopsis* sp3 | 0,2 | 0 | 0 | 0,8 | 0,2 |
|  | *Strumigenys* | *Strumigenys elongata* | 0 | 0 | 0 | 0,4 | 0 |
|  | *Temnothorax* | *Temnothorax subditivus* | 0 | 0 | 0 | 0 | 1 |
| Ponerinae | *Anochetus* | *Anochetus inermis* | 0 | 0 | 0,2 | 0,6 | 0,8 |
|  | *Leptogenys* | *Leptogenys pubeceps* | 0,2 | 0 | 0,8 | 0,8 | 0,2 |
|  |  | *Leptogenys volcanica* | 0 | 0 | 0 | 0,2 | 0 |
|  |  | *Leptogenys ritae* | 0 | 0,2 | 0,2 | 0,2 | 0 |
|  | *Neoponera* | *Neoponera villosa* | 0,2 | 0,6 | 0 | 0,4 | 0 |
|  | *Odontomachus* | *Odontomachus bauri* | 0 | 0,4 | 0,8 | 1 | 1 |
|  | *Pachycondyla* | *Pachycondyla harpax* | 0 | 0,2 | 0,8 | 0,6 | 0,6 |
|  |  | *Pachycondyla impressa* | 0,2 | 1 | 0,6 | 0,8 | 0 |
|  | *Platythyrea* | *Platythyrea pilosula* | 0 | 0 | 0 | 0,2 | 0 |
| Pseudomyrmecinae | *Pseudomyrmex* | *Pseudomyrmex* sp1 | 0,2 | 0 | 0 | 0,6 | 0,4 |
|  |  | *Pseudomyrmex boopis* | 0,4 | 0,6 | 0,8 | 0,8 | 0,8 |
|  |  | *Pseudomyrmex gracilis* | 0,2 | 0,6 | 0,2 | 0,8 | 0 |
|  |  | *Pseudomyrmex* sp4 | 0 | 0 | 0,2 | 0,2 | 0,2 |
|  |  | *Pseudomyrmex spinicola* | 0 | 0,2 | 0 | 0,4 | 0 |
|  |  | *Pseudomyrmex* sp6 | 0 | 0 | 0,2 | 0,2 | 0 |
|  |  | *Pseudomyrmex simplex* | 0 | 0,2 | 0,6 | 0,2 | 0 |
|  |  | *Pseudomyrmex* sp8 | 0 | 0,2 | 0 | 0,2 | 0 |
|  |  | *Pseudomyrmex venustus* | 0 | 0 | 0,6 | 0,2 | 0,4 |
|  |  | *Pseudomyrmex kuenckeli* | 0 | 0,4 | 0 | 0 | 0 |
|  |  | *Pseudomyrmex termitarius* | 0,2 | 0 | 0 | 0 | 0 |
|  |  | *Pseudomyrmex* sp11 | 0 | 0 | 0 | 0,2 | 0 |

The most species-rich genera included *Camponotus* Mayr, 1861 (13 species), *Pseudomyrmex* Mayr, 1861 (12), *Cephalotes* Latreille, 1802 (9), and *Pheidole* Westwood, 1839 (8) (**Table 3**).

The seven most frequently encountered species, accounting for 8.43% of total individuals, were *Pseudomyrmex boopis* (Roger, 1863), *Camponotus lindigi* Mayr, 1870, *Camponotus substitutus* Forel, 1899, *Solenopsis geminata* (Fabricius, 1804), *Cephalotes columbicus* (Forel, 1912), *Crematogaster ampla* Forel, 1912 and *Ectatomma ruidum* (Roger, 1860) (**Table 2**).

Among the sampling methods, Pitfall traps were the most effective in collecting a higher richness of morphospecies, genera, and subfamilies in our sites. Bait traps and manual collection showed intermediate effectiveness. The Malaise traps although they recorded lower relative values, contributed complementary records that enriched the sampled diversity. In total, seven species were captured exclusively by this trap, and in each site, between 2 and 12 unique species were recorded using this method (see figure S1 in the Supplementary Material)

We generated a comprehensive species list, incorporating records from literature, environmental reports, and this study. Here we identify eight subfamilies (Myrmicinae, Pseudomyrmecinae, Ponerinae, Amblyoponinae, Dolichoderinae, Ectatomminae, Dorylinae, and Formicinae), spanning 55 genera and 169 species (see Table S2 in the Supplementary Material)

### Alpha Diversity

Sampling completeness was generally high, exceeding 80% at most sites, except Los Charcones (70.19%). El Morro had the highest species richness (62 species), followed by Chorrera (43) and Uninorte (41). Libraria (31) and Los Charcones (31) exhibited the lowest richness (**Table 2; Table 3**). Ant community diversity varied significantly among sites (p = 5.3 × 10^−6^).

### Taxonomic Distinctness (TD)

Taxonomic Distinctness analysis (**Table 2; Figure 3**) showed that Los Charcones, Libraria, and El Morro fell within the expected range, with Libraria closest to the average. El Morro was slightly above the expected range, while Los Charcones was slightly below. Conversely, Uninorte and Chorrera were outside the model’s expected range, indicating significantly different taxonomic structures (p = 4.27 × 10^−3^, p = 3.63 × 10^−3^).

**Figure 1.**
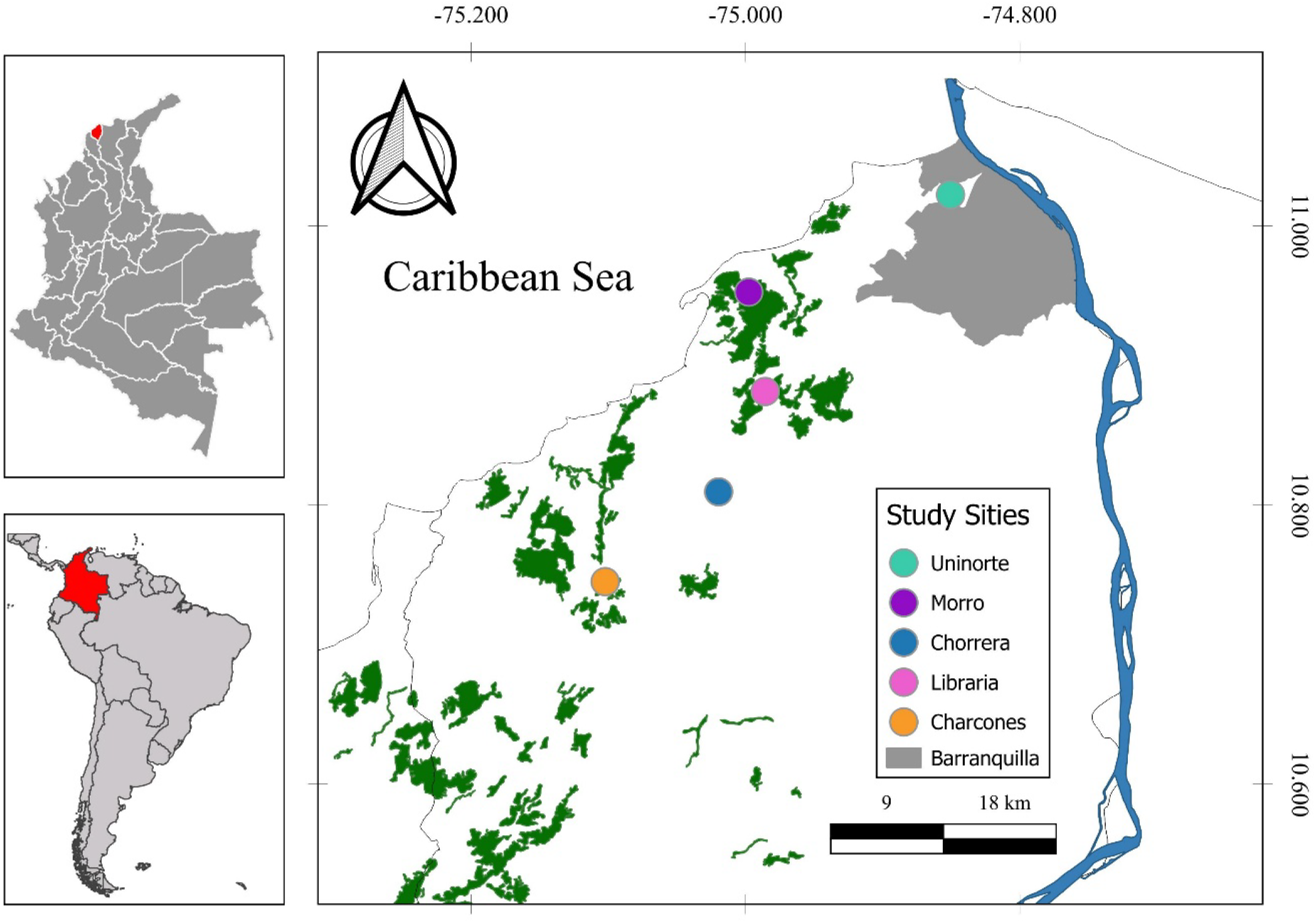
Study Area. The study was conducted at five sites with varying degrees of intervention. Uninorte, located within the ecocampus of the Universidad del Norte, corresponds to a fragment embedded within an urban matrix. El Morro is situated in an area fragmented by urban expansion processes. Libraría is a recovering fragment surrounded by areas dedicated to livestock and agriculture. Chorrera is a gallery forest within a civil reserve, with occasional cattle ingress. Los Charcones, also in a civil reserve, is a secondary forest that was previously used for livestock activities (41).

**Figure 2.**
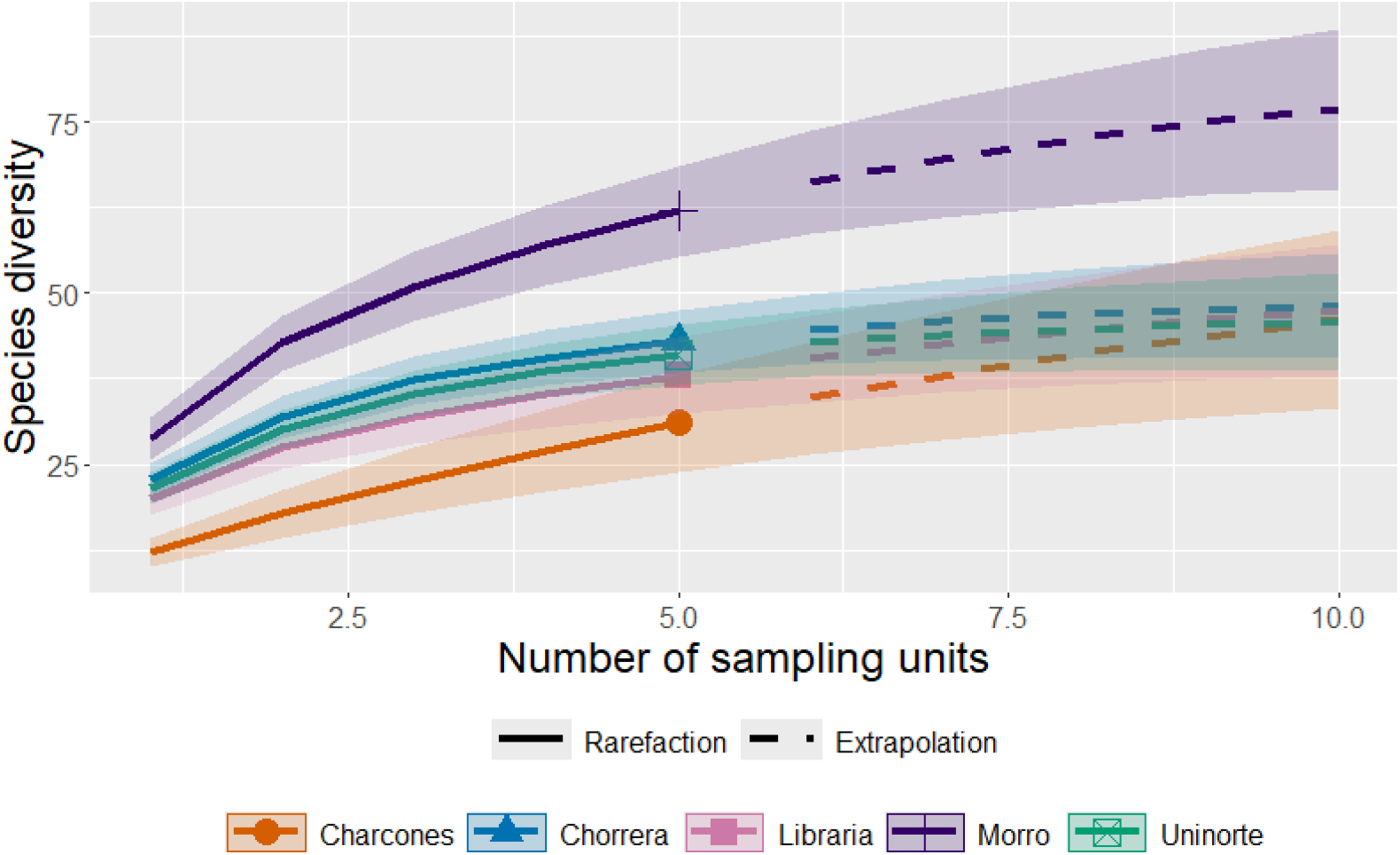
Rarefaction and extrapolation of Hill numbers, order q=0; Species count. Ant diversity showed significant differences among sites. El Morro was the most diverse, with 62 species, significantly surpassing the others. In contrast, Chorrera, Uninorte, Libraría, and Charcones did not show significant differences among each other, although their species richness ranged from 43 to 31 species.

**Figure 3.**
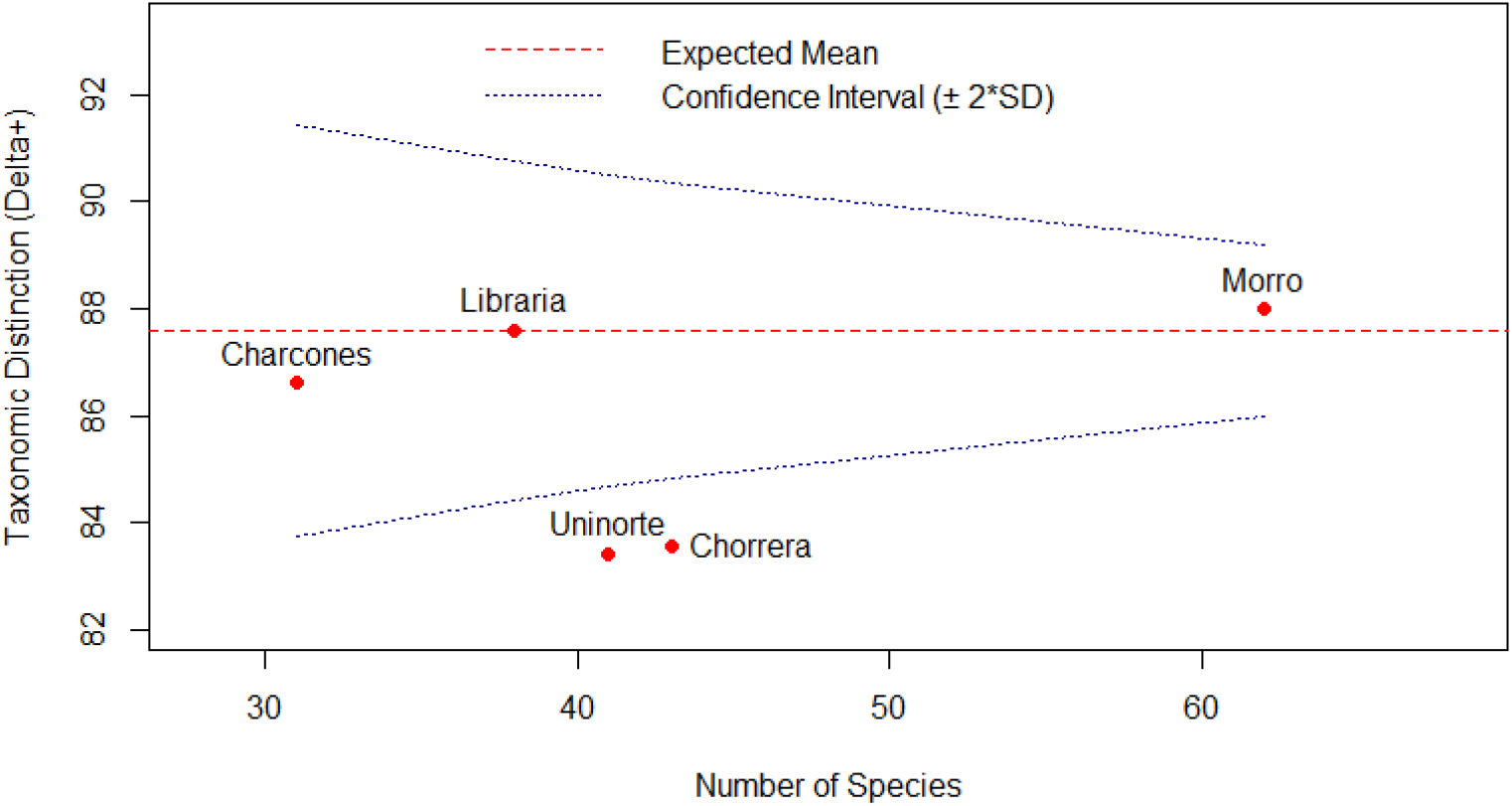
Funnel plot. The studied sites show clear differences in their phylogenetic diversity. El Morro has the highest species richness, and a higher taxonomic distinctness compared to the other sites, although it is not significantly different from the expected value. Libraría and Charcones have values close to the expected mean, suggesting communities with a balanced distribution across different lineages. In contrast, Uninorte and Chorrera exhibit significantly low taxonomic distinctness, indicating phylogenetically homogeneous communities dominated by species that are closely related.

### Beta Diversity

Of the 81 recorded morphospecies, 13 were present at all sites, including representatives from: *Camponotus* Mayr, 1861, *Cephalotes* Latreille, 1802, *Crematogaster* Lund, 1831, *Cyphomyrmex* Mayr, 1862, *Ectatomma* Smith, 1858, *Mycetomoellerius* Solomon et al., 2019, *Pheidole* Westwood, 1839, *Pseudomyrmex* Mayr, 1861, and *Solenopsis* Westwood, 1840 (**Table 4**).

**Table 4.** Shared and Unique Species at the Five Study Sites. Thirteen species were shared among the sites, while each site exhibited a set of unique species. El Morro was the site with the highest number of unique species, and it also recorded the presence of the invasive species *Tapinoma melanocephalum*.

| Shared Species | Los Charcones | Chorrera | Libreria | El Morro | Uninorte |
| --- | --- | --- | --- | --- | --- |
| *Camponotus lindigi* | *Camponotus dalmasi* | *Cephalotes atratus* | *Forelius damiani* | *Camponotus indianus* | *Nylanderia* sp1 |
| *Camponotus coruscus* | *Cephalotes pusillus* | *Pseudomyrmex kuenckeli* |  | *Camponotus linnaei* | *Camponotus* sp10 |
| *Cephalotes columbicus* | *Gnamptogenys striatula* | *Solenopsis* sp2 |  | *Dorymyrmex bicolor* | *Cephalotes* sp8 |
| *Cephalotes mompox* | *Pseudomyrmex termitarius* |  |  | *Ectatomma tuberculatum* | *Dorymyrmex biconis* |
| *Crematogaster ampla* |  |  |  | *Leptogenys volcanica* | *Rogeria foreli* |
| *Cyphomyrmex rimosus* |  |  |  | *Pheidole* sp4 | *Rogeria belti* |
| *Ectatomma ruidum* |  |  |  | *Pheidole* sp5 | *Temnothorax subditivus* |
| *Mycetomoellerius uruchii* |  |  |  | *Platythyrea pilosulla* |  |
| *Pheidole transversostriata* |  |  |  | *Pseudomyrmex* sp11 |  |
| *Pheidole* sp3 |  |  |  | *Sericomyrmex opacus* |  |
| *Pseudomyrmex boopis* |  |  |  | *Strumigenys elongata* |  |
| *Solenopsis geminata* |  |  |  | *Tapinoma melacephalum* |  |
| *Camponotus substitutus* |  |  |  |  |  |

Beta diversity analysis indicated that species turnover was the dominant factor in site-to-site variation, accounting for 59.1% of total dissimilarity, while nesting contributed only 11.8%. This suggests that diversity differences are likely explained by species replacement rather than a loss of species, but this should be thoroughly explored in subsequent studies.

Los Charcones exhibited the highest dissimilarity (56.25%–63.24%), likely due to its low species richness and limited shared species with other sites. Chorrera and El Morro had the lowest dissimilarity (43.18%), reflecting a high number of shared morphospecies. Uninorte showed the highest species turnover (39.22%–56.14%), followed by Los Charcones, Libraria, and Chorrera (**Figure 4**)

**Figure 4.**
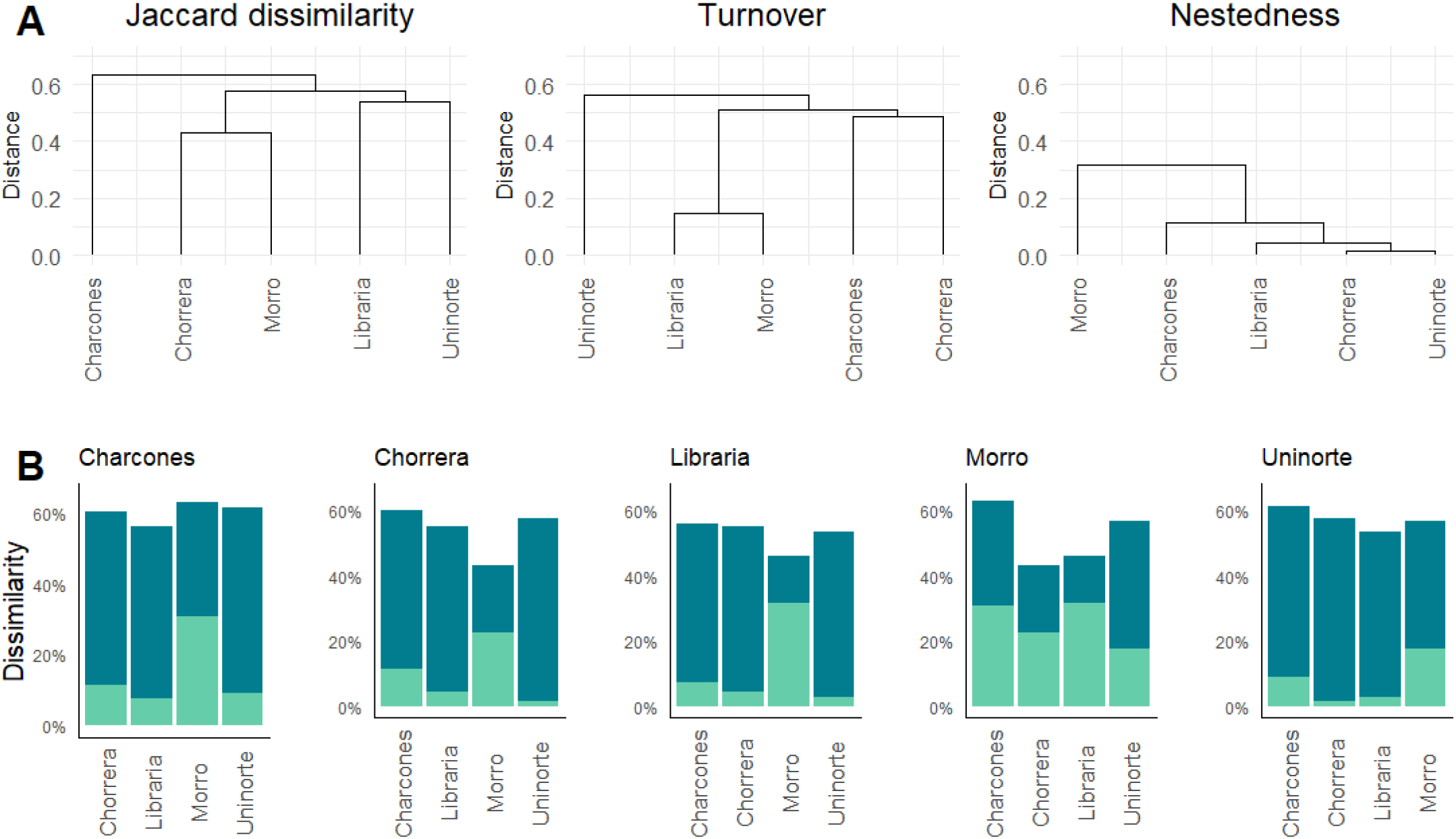
Beta Diversity. A) The dissimilarity in ant community composition was high (70%), primarily driven by species turnover (56.1%), with a smaller contribution from nestedness (11.8%). B) Turnover was especially marked in the Uninorte forest, indicating distinct species composition compared to the other sites. In contrast, nestedness was more prominent in El Morro, the most species-rich forest, suggesting that the other forests share a subset of its species

Despite low overall nesting, Los Charcones, Chorrera, Libraria, and Uninorte comprise an apparent subset of El Morro’s diversity, which is likely a local gradual loss of species relative to this site.

## 5. Discussion

This study provides the first taxonomic diversity analysis of all ants as a group in the department of Atlántico and presents an integrated species list for the region. The recorded species represent 6.75% of the ant species, 29.5% of the genera, and 63.3% of the subfamilies previously reported in Colombia (10). The species list derived from literature records encompasses 14% of the ant species, 52.4% of the genera, and 72.7% of the subfamilies documented in the country.

Among subfamilies, Myrmicinae was the most diverse, consistent with previous research in tropical dry forests (8,11–13,26). This richness is likely due to their diverse ecological habits (10,26,27). Formicinae was the second most diverse subfamily, mainly represented by *Camponotus* Mayr, 1861, known for its adaptability to disturbed and varied environments (27). Pseudomyrmecinae, represented by *Pseudomyrmex* Mayr, 1861, specializes in arboreal habitats and frequently establishes symbiotic relationships with plants such as acacias (10,27).

Our study documented 87.5% of the subfamilies previously reported in the department’s tropical dry forests, with Amblyoponinae—reported by Domínguez-Haydar et al. (2007) (11)—absent from our samples. Additionally, we identified 60.63% of the 55 genera previously recorded for the department (11–13,16,17,28). Notably, we report the presence of the genus *Forelius* Emery, 1888 for the first time in Atlántico.

Although Malaise traps are not commonly used for ant collection, in this study these traps captured 40 morphospecies, with an average of 17 per site. In previous studies that used a similar set of traps but included Winkler traps instead of Malaise traps, fewer species were recorded per site (28 and 26 morphospecies, respectively) (12). Malaise traps provide a valuable complement to pitfall or Winkler traps, which primarily collect ground- and leaf-litter-dwelling ants, by capturing species that do not dwell on the ground or in the leaf litter but rather inhabit branches and the canopy and therefore would not be detected by these methods.

### Distribution Expansions

Our findings extend the known distribution of 16 species within Atlántico, including *Azteca forelii* Emery, 1893, *Camponotus bidens* Mayr, 1870, *Camponotus indianus* Forel, 1879, *Camponotus linaeii* Forel, 1886, *Crematogaster carinata* Mayr, 1862, *Dorymyrmex bicolor* Wheeler, 1906, *Forelius damiani* Guerrero & Fernández, 2008, *Leptogenys pubeceps* Emery, 1890, *Leptogenys volcanica* Lattke, 2011, *Leptogenys ritae* Forel, 1899, *Paratrachymyrmex irmgardae* (Forel, 1912), *Pheidole transversostriata* Mayr, 1887, *Pseudomyrmex kuncklei* (Emery, 1890), *Pseudomyrmex termitarius* (Smith, 1855), *Rogeria belti* Mann, 1922, and *Temnothorax subditivus* (Wheeler, 1903).

Additionally, we provide the first record of *Leptogenys volcanica* Lattke, 2011 in the Colombian Caribbean region, expanding its known distribution beyond Valle del Cauca and Panamá (10,17). This suggests a broader range for the species and raises the possibility of its presence in other Caribbean departments.

### Taxonomic Distinction and Habitat Differences

The taxonomic distinctness (TD) values suggest important ecological differences among sites. El Morro exhibited the highest TD value, likely due to its greater species, genus, and subfamily richness. Chorrera and Uninorte, despite sharing the same number of subfamilies as El Morro, had lower TD values, with Myrmicinae dominating their assemblages (60.47% and 58.54%, respectively).

El Morro’s high taxonomic distinction and diversity suggests a structurally complex ecosystem supporting multiple ecological niches (29). Similar patterns have been observed in tropical dry forests for other biological groups, such as birds, where both taxonomic and functional diversity increase as forests mature (30).

In contrast, Los Charcones exhibited lower arboreal genus representation (e.g., *Cephalotes, Pseudomyrmex*) than other sites, likely due to past anthropogenic disturbances such as livestock farming 12 years ago (31). Land-use changes disrupt forest flora and fauna, altering ecological guilds and delaying recovery for decades (9).

The Uninorte fragment had the lowest TD value, likely due to its isolation within an urban matrix lacking connectivity with other forests. This restricts species colonization and limits local biodiversity (32).

### Community Composition and Fragmentation

Our results indicate that each forest fragment harbors a unique ant community, with high species turnover between sites. Environmental characteristics likely shape these differences (33,34). Despite this variability, 13 species were present in all fragments, representing widely distributed genera that thrive in disturbed forests (35–37).

Conversely, many fragment-specific morphospecies belong to cryptic genera such as *Strumigenys* Smith, 1860, *Sericomyrmex* Mayr, 1865, *Leptogenys* Roger, 1861, *Temnothorax* Mayr, 1861, *Rogeria* Emery, 1894, *Gnamptogenys* Roger, 1863, *Brachymyrmex* Mayr, 1868, and *Forelius* Emery, 188. These species contribute significantly to community heterogeneity and are particularly sensitive to habitat disturbance (10,38).

### Nesting and Fragment Roles

El Morro exhibited the highest nesting values, suggesting that many species recorded in other fragments form a subset of its diversity. This indicates that species loss in other fragments may be linked to land-use changes such as urbanization, agriculture, and livestock farming. The high taxonomic distinctness and presence of exclusive, cryptic species reinforce the idea that El Morro provides more favorable conditions for biodiversity. These results align with previous studies in tropical dry forests linking high nesting values to well-preserved ecosystems (9,36).

In contrast, Uninorte exhibited the highest species turnover, likely due to its isolation within an urban landscape (39). This suggests that Uninorte might function as a diversity sink, with limited species exchange with other fragments, whereas El Morro could act either as a diversity source or a representative of less altered diversity. Similar patterns have been observed in Caribbean tropical dry forests, where fragmentation and isolation have created diversity sinks for various biological groups, such as dung beetles (40).

## 6. Conclusion

This study provides a comprehensive assessment of ant diversity in tropical dry forest fragments of Atlántico, expanding species distribution records and demonstrating that habitat fragmentation and land-use change significantly impact both taxonomic and phylogenetic diversity. Our results support the hypothesis that well-preserved fragments, such as El Morro, harbor more diverse and representative evolutionary lineages, highlighting their ecological value as biodiversity reservoirs. Future conservation strategies should prioritize habitat connectivity to reduce biodiversity loss in fragmented landscapes.

